# Bursts of novel composite gene families at major nodes in animal evolution

**DOI:** 10.1101/2023.07.10.548381

**Authors:** Peter O. Mulhair, Raymond J. Moran, Jananan S. Pathmanathan, Duncan Sussfeld, Christopher J. Creevey, Karen Siu-Ting, Fiona J. Whelan, Davide Pisani, Bede Constantinides, Eric Pelletier, Philippe Lopez, Eric Bapteste, James O. McInerney, Mary J. O’Connell

## Abstract

A molecular level perspective on how novel phenotypes evolve is contingent on our understanding of how genomes evolve through time, and of particular interest is how novel elements emerge or are lost. Mechanisms of protein evolution such as gene duplication have been well established. Studies of gene fusion events show they often generate novel functions and adaptive benefits. Identifying gene fusion and fission events on a genome scale allows us to establish the mode and tempo of emergence of composite genes across the animal tree of life, and allows us to test the repeatability of evolution in terms of determining how often composite genes can arise independently. Here we show that ∼5% of all animal gene families are composite, and their phylogenetic distribution suggests an abrupt, rather than gradual, emergence during animal evolution. We find that gene fusion occurs at a higher rate than fission (73.3% vs 25.4%) in animal composite genes, but many gene fusions (79% of the 73.3%) have more complex patterns including subsequent fission or loss. We demonstrate that nodes such as Bilateria, Euteleostomi, and Eutheria, have significantly higher rates of accumulation of composite genes. We observe that in general deuterostomes have a greater amount of composite genes as compared to protostomes. Intriguingly, up to 41% of composite gene families have evolved independently in different clades showing that the same solutions to protein innovation have evolved time and again in animals.

**Significance statement:** New genes emerge and are lost from genomes over time. Mechanisms that can produce new genes include, but are not limited to, gene duplication, retrotransposition, *de novo* gene genesis, and gene fusion/fission. In this work, we show that new genes formed by fusing distinct homologous gene families together comprise a significant portion of the animal proteome. Their pattern of emergence through time is not gradual throughout the animal phylogeny - it is intensified on nodes of major transition in animal phylogeny. Interestingly, we see that evolution replays the tape frequently in these genes with 41% of gene fusion/fission events occurring independently throughout animal evolution.

## Introduction

Composite genes emerge by fusion of distinct protein coding sequences (“components”), or by the fission of protein coding sequences into components. Often composite genes establish novel domain architectures, expression profiles, and functions (1–7). For example, the fusion gene *Jingwei* is remodelled from *yellow emperor* and *alcohol dehydrogenase* genes combining activity on both long chain alcohols and diols, including growth hormones and pheromones, and establishing a novel developmental function in Drosophila (8). In addition, the *kua-UEV* fusion gene in human has facilitated cytoplasmic localization of an otherwise solely nuclear polyubiquitination co-effector (9). Whilst there is mounting evidence for the role of gene fusion in driving adaptive evolution (10–15), there are a number of outstanding questions about the evolution of composite genes in animals. Specifically, how prevalent composite gene formation has been, whether the emergence of composite genes occurs in bursts rather than gradually, whether the pattern of emergence of composite genes correlates with the origin of major animal groups, and at what rate fusions or fissions occur. In addition, whilst the convergent evolution of phenotypes is well established in animals, the extent to which composite genes can arise independently in different lineages is largely unknown. Given the divergence in morphologies, niches, lifestyles, and indeed genomes, it is not clear whether repeated evolution of the same molecular components would be precluded, or whether the deterministic effects, *i.e.* the benefits of particular kinds of composites, would overcome any contingent effects of prior genome evolution.

Animal genome evolution has been shaped by regulatory innovations (16), by *de novo* gene genesis (17–21), and by gene duplication and gene loss (20, 22). Composite genes have been particularly challenging to study as they simultaneously reside in more than one homologous gene cluster which complicates gene family assignment and phylogenetic analyses. Studies of protein coding aspects of animal evolution have necessarily relied upon strict definitions of gene families that limit our view to purely furcating processes (23, 24). An alternative view is provided by retaining the connections between composite genes and their components in sequence similarity networks (SSNs), permitting genes to be members of more than one family simultaneously. Taking advantage of the unique network motif typical of composite genes, *i.e*. they form “non-transitive triplets” thereby connecting gene families that are otherwise unconnected through partial sequence homology, we can identify candidate composite genes from genome scale data (25).

The protein domain space that comprises all animal proteins (components and composites alike) is limited, and it is conceivable that the same composite gene forming event could occur multiple times independently (26). Indeed, comparative empirical studies reveal surprisingly repeatable evolutionary fates in closely related lineages, a trend considered to reduce with increasing distance (27, 28). To date, the rate of independent evolution observed within multi-domain proteins has varied dramatically from very low, *i.e.* 0.4% (29), to much higher, 5-25% (13, 27). The lower range of estimates of independent evolution is thought to be caused by limited taxon sampling (13, 27). Using a large representative dataset, we provide a statistically sound framework to elucidate the rate of independent evolution of composite genes in animals.

## Materials and methods

### Dataset assembly

A dataset of 1,217,174 protein coding genes from a sample of 63 animal species representing all major clades within the animal tree was obtained from the OMA orthology database (70). Taxa were sampled to capture the known periods of major transition within animal evolution, and species representing all major nodes in the animal tree were included. The quality of data used was of particular importance in this study (given the potential for misidentification of composite genes) therefore taxon sampling was guided by the quality of gene annotation of the available species genomes using two filtering steps of genomes. First, we searched for protein coding genes known to be present across all of Metazoa (412 genes in total) (71), ranking genomes as high quality if they possessed >70% of the conserved set, while low quality genomes had less. Next, a smaller set of 40 protein coding genes that are annotated as being present across all of life (72) were used as queries to search for their presence in the set of animal genomes. As this set of protein coding genes is more conserved, this allowed for stricter filtering for quality of the genomes. All homology searching for the “core set of metazoans” and “all of life” protein coding genes was carried out using a reciprocal BLASTp approach (73). Searching for the set of conserved genes within sampled genomes in Metazoa and all of life, ensured that genomes of high quality (deemed by the presence of these sets of conserved genes) were used in our analysis.

To construct a time-calibrated species tree, node dates and topology were obtained from TimeTree (53), and contentious groupings (such as the branching order at the root of the animal lineage) were resolved based on current literature on the animal phylogeny (50–52, 74). We also included an alternative topology for the root position on the animal tree to test if our results are robust to the position of the deepest divergences. Twelve of the species in our dataset were missing from the TimeTree database, and so to place their position and time of divergence, closely related species to these lineages were used as replacements. In most cases, sister species from the same genus were present, and a list of the closely related species used to replace them can be found in (**Supplementary Table S2**). With other species, such as the case of *Ciona savignyi,* which was not present in TimeTree, the divergence time between it and its sister lineage *Ciona intestinalis* was taken as 176 MYA from the literature (75).

### Generation and filtering of the sequence similarity network

An all versus all BLASTp (Altschul et al. 1990) was carried out (E-value <= 1e-5, percent identity >= 30%). The statements of homology output from BLASTp were used to generate a Sequence Similarity Network (SSN), using the cleanBlastp step in CompositeSearch (25). CompositeSearch applies a modified Depth First Search (DFS) algorithm to annotate gene families followed by subsequent network searching to define composite genes and gene families, then takes this SSN as input and identifies composite gene clusters which are denoted by non-transitive triplet patterns in the SSN. We used an E-value cutoff of 1e-5, percent identity cutoff of 30%, and coverage threshold of 80%. This provides output files on all HGs - both the gene families detected, and the gene families annotated as composite (CHGs). The composite gene family’s annotation file also provides information on the size of the composite gene families, the number of component families associated with the composite family, the size of the component gene families and the connectivity of the subgraph of composite genes within a family. Information such as the number of composite genes within a family, and the amount of overlap of the homologous regions between the distinct component genes and the composite gene is all made readily available. As discussed previously the detection of composite genes may be prone to misidentification and false positives. Therefore, as an initial filtering step, a series of quality filters were applied to the putative composite gene families identified in the CompositeSearch analysis: firstly, singleton CHGs, *i.e.* those with only a single member in the CHG, were removed. In total this filtering step removed 48,640 CHGs out of the total of 77,085 putative CHGs. In addition, genes where the mapping of components to the composite was ambiguous or where the mapping of the components overlapped, were also removed (this removed a further 14,813), leaving a total of 13,632 remaining putative CHGs.

### Finding evidence for expression of unique joining-points of composite genes using publicly available transcriptome data

We collated a dataset of all available transcriptomes from RNA sequencing studies for each taxon (52 out of 63 taxa had RNAseq data available). This allowed us to assess the putative composite genes at two levels: (i) their validity - making sure we do not report putative composite genes that are misassembly or misannotation artefacts, and (ii) whether they have evidence of expression. RNAseq reads were mapped to the unique joining-point region of the composite genes (*i.e.* the junction of component genes) using bowtie2 (76). RNAseq datasets were selected based on their robustness (as measured by the number of time points and tissues sampled) and the phylogenetic distribution of composites. For example, for widely distributed composite families, representative taxa from across the lineages containing the composite family were chosen based on the robustness of their available RNAseq datasets. The representative taxa for each of the Bilaterian clades included humans (Deuterostomia) and fruit-fly (Protostomia). Coverage across the composite joining- point was assessed using BEDTools (77). Evidence for transcription of the composite gene was determined by the coverage of at least one read across the joining-point.

### Domain architecture analysis

For all of our HG datasets (composite HGs, component HGs, and non-composite associated HGs) we first annotated protein domains from the Pfam database using domain-specific hidden markov models (31), using pfam_scan.pl and parsing using PfamScanner with an e-value threshold of 1e-3. We calculated the proportion of retention and loss during gene fusion for each domain by dividing every time it is present or lost in a composite gene by the number of times it is seen in all component genes (this analysis was carried out on just the domain type architecture of the proteins rather than the full protein architecture which may include repeat domains). No statistically significant correlation is observed between the number of times a domain type is seen in a CHG vs the proportion of time it is present or lost (**Supplementary Figure S1**). Similarly, when we annotate the domain by function or size, there does not seem to be a correlation between the presence of a domain and these traits.

To assess whether there were any domains enriched in the set of composite genes which were annotated as emerging independently multiple times, we compared domain sets between single event CHGs and convergently formed CHGs. We obtained domain lists and correlating functions for both sets of CHGs and applied the find_enrichment.py script from Goatools (78) setting the full set of domains from all CHGs as the background.

### Assessment of rate of convergent evolution of CHGs

The pipeline for determining which CHGs emerged in a single event or multiple events is highlighted in **Figure 3**. In summary, we first retained all CHGs that had a simple 2-to-1 relationship between composite and component genes. Next, we extracted the homologous regions between both parts of the composite gene and their respective component sequences, using information from the tabular all-v-all BLAST output. The homologous regions between composite and components were aligned using MAFFT (79) and trimmed using trimAl, using the -gappyout parameter (80). Corresponding gene trees for both parts of the composite gene and respective component genes were constructed using IQTree (42, 43), applying ModelFinder (81) to find the model of best fit and carrying out 1000 ultrafast bootstrap replicates. We used clan-check to classify composite genes according to whether they appeared at face-value to have a single or multiple origins (82). Next, we constructed constrained gene trees in IQTree by forcing the composite sequences to be monophyletic and applying the model of best fit as inferred from the previous gene tree construction step. This ensures that gene tree construction is consistent between the two approaches, the only exception being that the composite genes are forced to be monophyletic. Finally, an AU test was carried out using IQTree, applying the -au parameter to compare support levels for the inferred gene tree with the constrained gene tree.

We also assessed the conservation of the joining-points between composite genes in each CHG tested. This involved determining the location of the joining-point for each composite gene by annotating where the sequence homology of the component genes mapped to the composite gene. Then, each composite gene in each CHG was split into four non overlapping but equal length regions (proportional to overall length of the composites) and we assessed whether the joining-points for all composite genes in a CHG fell within the same region. The assumption being that, while there may be some variation in the exact location of the joining-point, those in the same region of the composite gene provide more support for a single origin of a given CHG. This test was carried out on all CHGs. Leading on from this we could then address the question of whether we observe different joining-points in CHGs of multiple origin.

### Mapping composite gene gain and loss onto species tree

For the taxa in our dataset, most of the branching patterns are well resolved allowing the analysis of the rate of emergence of CHGs on each branch across the tree to determine the patterns of gain and loss. We reconstructed the gain/loss history of the CHGs and used a constrained timetree (53) to determine their rates of gain and loss. The pattern of gains and losses of CHGs across the tree was assessed using one of two models; if CHGs were determined to have been formed in a single event, we used Claddis (46) an R package which operates in a maximum likelihood framework to describe characteristics of binary data. Specifically we used the map_dollo_changes.R function, which was developed to generate a stochastic character map for Dollo characters, allowing for a single gain event followed by any number of losses (47). Alternatively, if CHGs were annotated as evolving convergently, we implemented the Mk model using a stochastic character mapping approach, this time implemented in RevBayes (49). Setting the root to zero and using a Mk model with unequal transition rates, allowing a character to be lost and gained a number of times at different rates across the tree, we measured the rate of gain and loss of each CHG individually. For each CHG we ran two mcmc chains for 5,000 generations each, allowing us to measure the precise timing of gain along each branch stochastically. Visualisation of the numbers of gains and losses on the species tree was carried out using an edited version of the R package, RevGadgets (83). To determine the rate of gain and loss of the CHGs mapped to the species phylogeny, we divided the number of CHGs gained or lost at a given node by the age of that node. The rates of gain and loss were added to each branch in the tree using ETE3 (84) and the tree was plotted using ggtree (85).

### Characterising composite formation events

Composite genes may have originated from either fusion or fission. Fusion events merge two or more pre-existing components (*e.g.* gene families without direct connections in sequence similarity networks). Fission results in the subsequent appearance of split forms (*i.e.* components) of the gene. CompositeSearch (25) does not natively provide a classification of detected composites as corresponding to a fusion or fission event, yet this step is pivotal for a deeper biological interpretation of the computational outcome. We therefore applied a phylogeny-based method to infer the relationship of evolutionary precedence between composites and their respective components, and deduce the type of gene remodelling that was detected in the network (*i.e.* the evolution of composite genes by fusion vs the evolution of component genes by fission).

The last common ancestor of each CHG and the last common ancestor of each of its components were mapped onto the reference species tree of our sample set. These last common ancestors represent the putative points of appearance (assuming a unique origin) of each composite family and their associated component families. A simple heuristic was then applied to label CHGs as fused (originated from a gene fusion event) or split (underwent a gene fission event). CHGs for which components existed prior to the composite origin were considered as fused, as the inference of a component evolutionarily older than the composite indicates that at least one “building block” of the composite was present in its ancestral lineage before its appearance, and thus was unlikely the result of the composite fission. Conversely, CHGs for which components appeared only below the composite origin were marked as having undergone gene fission, as split forms of the composite persist in extant lineages ancestrally carrying the non-split gene. Many CHGs exhibited a particular pattern with both a component existing prior to the composite form and another component evolving only after the composite origin. Such cases are difficult to ascribe to a single fusion or fission event and may be the result of a more complex evolutionary path: the existence of a component predating the CHG indicates that it likely originated from a fusion event, possibly followed by a subsequent gene fission or loss that would have given rise to the later-evolving component.

### Functional annotation of composite genes

All proteins used in our starting dataset were functionally annotated using eggNOG-mapper (86) v2.1.6, employing DIAMOND v2.0.11 to align sequences to the eggNOG database v5.0.2. Cluster of Orthologous Groups (COG) functional categories were extracted for each sequence, and where required these were used to annotate the representative set of functions per CHG. To first test whether there were differences between the functional groups represented in fusion genes versus genes never associated with CHGs, we compared each COG category for each gene annotated as fusion or annotated as neither fusion or fission, and plotted the relative proportions (*i.e.* the number of COG categories divided by the number of genes) for each category. Next, to compare proportion of functional categories which emerged on each node of the tree, including major animal nodes, we took the proportions of categories from each CHG, and inferred the overall contribution at each node by dividing all the categories by the number of CHGs gained on the given node. These proportions for each node were plotted individually for the major nodes and combined to compare against all other internal nodes in the tree.

## Results

### Sequence similarity networks uncover a large number of composite genes

From a set of 1.2 million protein coding sequences across 63 animal genomes we identified 297,806 homologous groups (HGs) of which 77,085 contained putative composite homologous groups (CHGs). We removed (i) all singleton CHGs (of which there were 48,640), and (ii) all putative CHGs where the contributing component sequences did not map to specific and non-overlapping regions of the composite gene (14,813 CHGs in total). Under these strict criteria we identified 13,632 CHGs, or ∼5% of all the gene families in animals, and these groups included 157,206 individual composite genes. To further mitigate against annotation and assembly artefacts, we assessed whether putative composites have associated evidence of gene expression. We mapped the unique “joining-point” in each composite gene to available transcriptome datasets (the “joining-point” is the location within the composite sequence where the contributing component sequences meet). Transcriptome data was available for 52 of the 63 species and 12,048 of the 13,632 CHGs (see Materials and Methods). A total of 7,774 CHGs (65%) had evidence of expression for at least one composite gene member of a given CHG family. The proportion with evidence of expression (*i.e.* 65%) is what we might expect from large scale RNAseq studies on temporal and spatial variation in expression in animal protein coding genes (30).

The 13,632 CHGs were related to a total of 40,217 component HGs, with the majority of CHGs (*i.e.* 10,855 (80%)) having just 2 component genes (or parts thereof). We identified a nested characteristic of composite formation in that CHGs once formed tend to contribute to further composite formation. In total, 11,805 out of 13,632 (87%) CHGs are also components (**Figure 1 a&b**). Indeed, when compared to genes that did not arise by composite formation, genes that arose by composite formation are more likely to subsequently contribute to other novel composite events (on average 10 vs 17 subsequent events respectively), suggesting there is a pool of genes prone to remodelling in animal genomes. On assessing protein length and domain content across the ∼1.2 million protein coding genes in the network we observe that composite genes display a wider range of domain combinations (as classified by Pfam (31)) than either (i) component genes (p < 2.2e-16, Wilcoxon signed-rank test), or (ii) non-composite associated genes (*i.e.* those genes that are not composite and not component) (p < 2.2e-16). Additionally, composite genes tend to have longer protein coding sequences than either component genes (p < 2.2e-16) or non-composite associated genes (p < 2.2e-16). On average component genes contribute 100 amino acids during composite formation - corresponding closely to the average length of a domain in all component genes in our dataset (*i.e.* 118 amino acids). The most common proportion of component gene sequence to be present following a fusion event is 20% or 100%, suggesting that the domain unit places the strongest constraint on the size and architecture of composite genes (**Figure 1c**). Comparing domain types between component and composite genes, we find that no domain is significantly over-represented across all CHGs and thus present at a higher rate than others during composite formation (27). However, whilst not statistically significant, domains WD40 and zf-C2H2 are present at a higher rate in CHGs than any other domain, *i.e.* WD40 is present in 391 components and is present in the resulting CHGs 58% of the time, and zf-C2H2 is present in 339 components and present in CHGs 58% of the time (**Supplementary Figure S1**). This trend possibly reflects the abundance and promiscuity of these two domains: the WD40 domain is one of the most abundant and also amongst the top interacting domains in eukaryotic genome (32), whilst the zf-C2H2 domain is amongst the most numerous of domains in metazoa (33).

**Figure 1.**
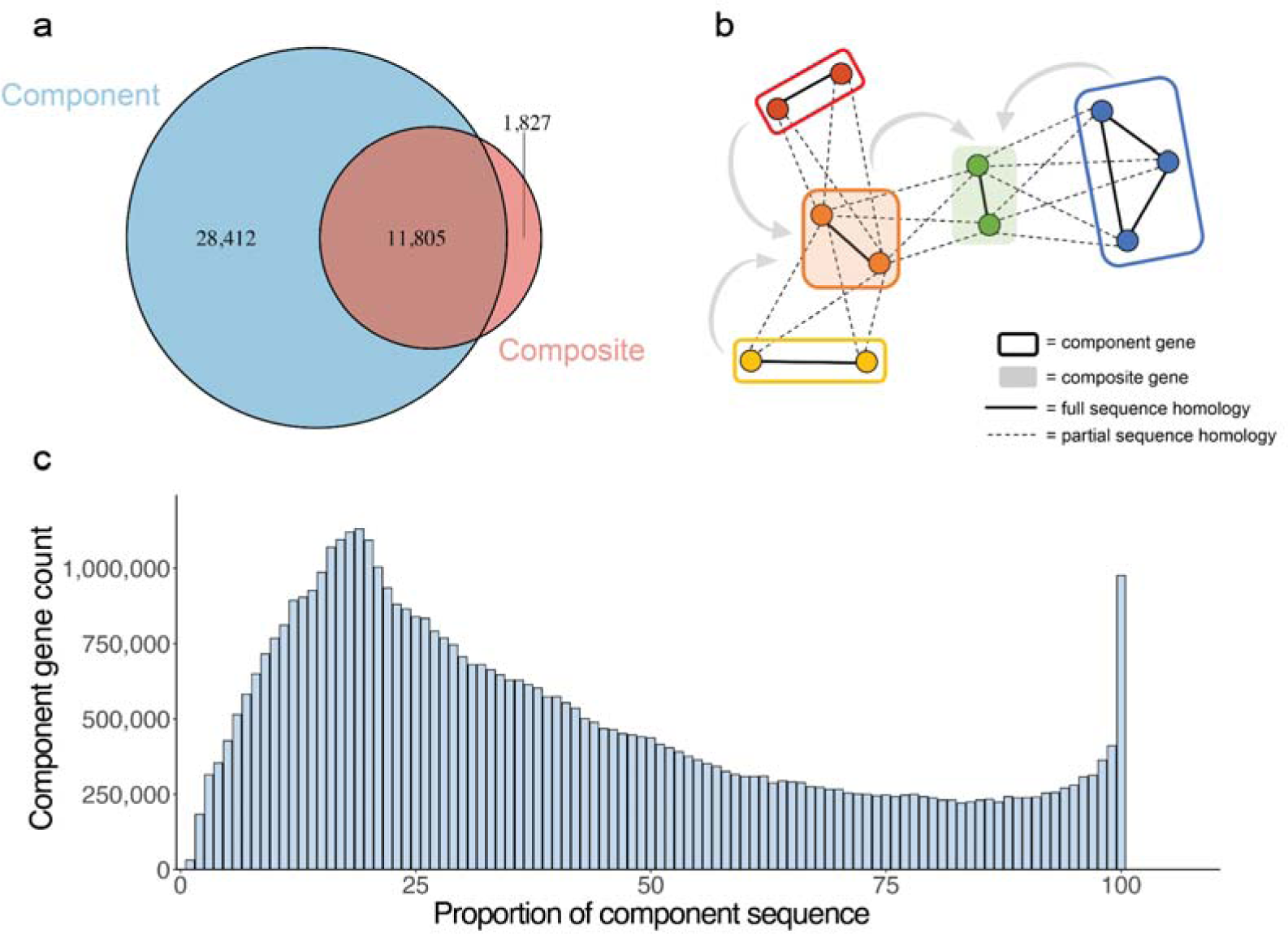
Characterisation and contributions of composite genes and their components. **(a)** The number of component CHGs (blue) and composite CHGs (red). Overlap represents component CHGs which are themselves composite. **(b)** Cartoon network demonstrating the nested nature of composite formation, whereby *e.g.* a CHG (orange) formed from distinct component CHGs in red and yellow, may itself be involved in a separate fusion with another HG component family (in blue) to form a new CHG (green). **(c)** Distribution of the component sequences showing the proportion of the component that is present in the composite.

For most CHGs (82% or 11,211 of 13,632 CHGs), the contributing elements (*i.e.* the parents of the gene fusion or the gene parent for the fission), are lost from the host genome. There are 2,421/13,632 CHGs where the composite and at least one component reside in the same genome simultaneously. For example, whilst previous studies of insulin-like growth factor-binding protein gene family (IGFBP) have characterised the functional domains, our analyses identify that the formation of this gene was via gene fusion on the stem chordate lineage (**Figure 2a**). We also show that following formation of this gene fusion its component genes were not lost from the genome. The process of gene fusion involved the C terminal IGFBP domain which functions to regulate IGF, and the N terminal Thyroglobulin-1 domain which contains nuclear localization sequences (**Figure 2b**). The IGFBP fusion and subsequent duplication resulted in many novel IGF-independent actions in a new cellular functional landscape (34–38). This example provides new insights into the process of gene fusion which involves retention of both component and composite genes in most chordates sampled (**Figure 2**). Conversely, the Nitrilase and fragile histidine triad fusion protein (NitFhit) demonstrates the loss of ancestral components following gene fusion. Given that the separate components have been found to be expressed and localised at similar time points and are also involved in similar interaction networks and functions, this fusion represents a coordination of biochemical pathways (39, 40). The NitFhit fusion was initially proposed to have originated by gene fusion in *C. elegans* and *D. melanogaster* (39), and we identify it in both Ecdysozoa and Lophotrochoza, placing the origin of the NitFhit fusion at the base of the Protostomia.

**Figure 2.**
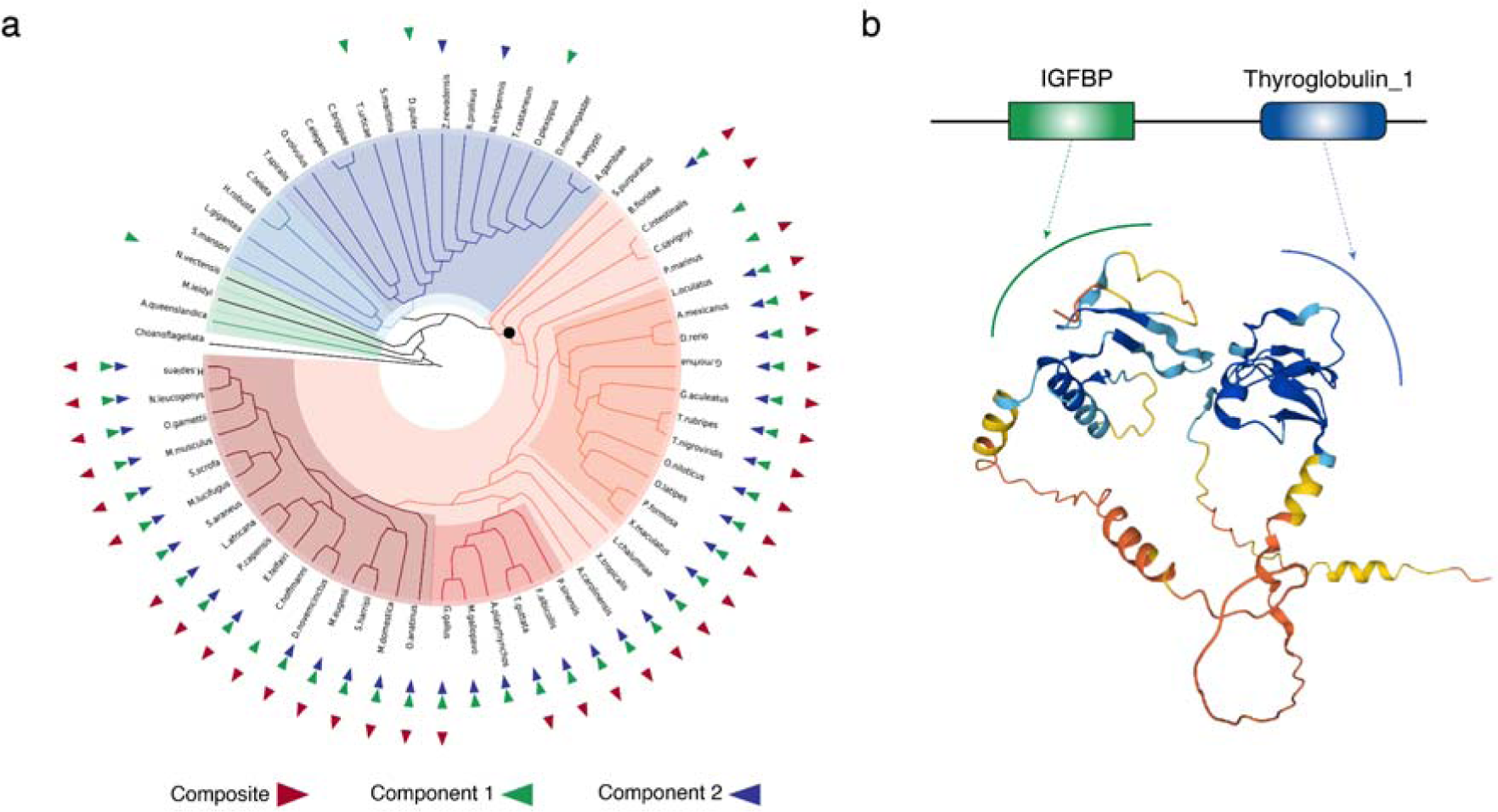
Evolution of the composite IGFBP gene in Chordates. **(a)** The chordate clade is highlighted with a red toned background on the circular tree, and the outgroups in blue and green toned backgrounds. The presence/absence of the composite and components are denoted with coloured triangles on the leaf nodes: the presence of the IGFBP composite gene is denoted as a red triangle, and the two component genes IGFBP and Thyroglobulin 1 are denoted as blue and green triangles respectively. The node of origin of the IGFBP composite genes is annotated by an in-filled circle on the species tree. **(b)** A cartoon of the constituent domains IGFBP (green) and Thyroglobin_1 (blue). Arrows point to the corresponding regions in the IGFBP composite gene protein structure. The structure colour gradient represents regions of high (blue) and low (red) confidence, note the two component protein domains are linked by a sequence of low structural confidence.

**Figure 3.**
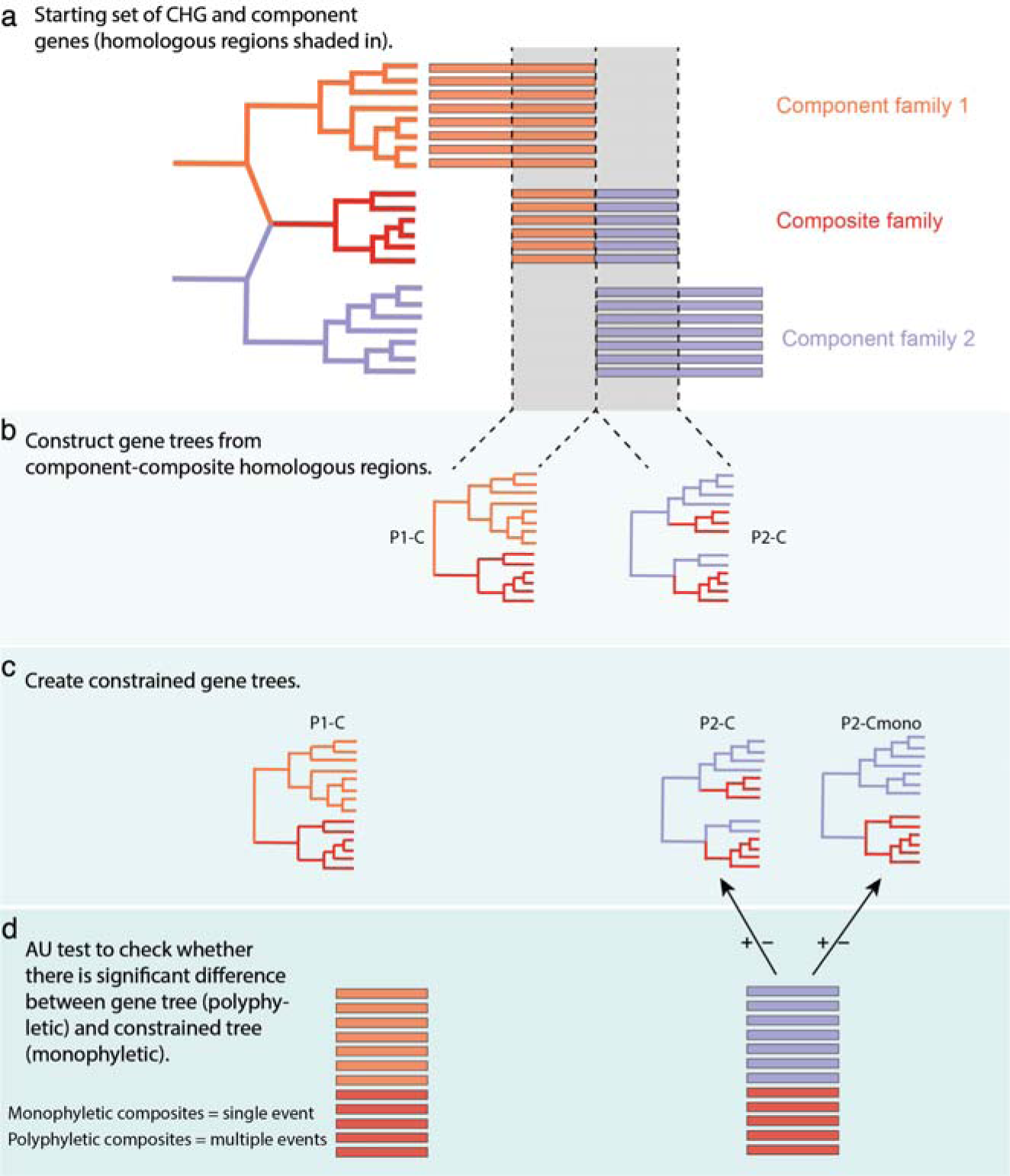
Pipeline used to identify composite genes that emerged in a single event or multiple independent events. **(a)** Summary of component and composite homologous sequences used for gene tree inference. **(b)** Gene trees inferred from component-composite sequences showing an example where the composite sequences (in red) form a monophyletic group (left) and an example where they do not (right). **(c)** Constrained trees inferred using the topology of the gene tree inferred from the previous step but forcing the composite sequences to be monophyletic. **(d)** Approximately unbiased test: measuring the significance in the difference in support for the constraint tree (monophyletic) versus the unconstrained (polyphyletic) tree. Where *H_0_* = no significant difference in likelihood score for the constraint and unconstrained trees.

### Larger number of composite genes are formed by gene fusion

As composite gene losses may be conflated with lineage-specific gene fission following a fusion event, we further categorised the mode of origin of all composite genes in each CHG as having emerged by gene fusion or fission (see Materials and methods, **Supplementary Figure S2**). Briefly, if the last common ancestor of the component genes was an older node than that of the composite gene, the composite was categorised as formed by gene fusion; and the converse for fissions. Out of the 13,632 CHGs in this analysis, 9,994 CHGs (73.3%) were found to have originated from a fusion event. Of these, 2,096 (21%) fit the scenario of a single fusion event with no subsequent fission, while 7,898 (79%) were inferred to have undergone a more complex pattern of independent fusion and/or subsequent fission events (**Supplementary Figure S2B**). Interestingly, there were 3,460 CHGs (25.4%) that underwent a single, unique fission event (**Supplementary Figure S2C)**. Finally, 178 CHGs (1.3%) could not be assigned to any of the above categories, suggesting a more complex evolutionary history. Overall, the relative rates of fusion and fission observed mirror recent findings which suggest that gene fusion played a greater role in metazoan gene family evolution as compared to other eukaryotes, *i.e.* Fungi (21, 41) (**Supplementary Figure S2)**. These findings also suggest that gene fission occurs at a previously underestimated rate in animal genomes, particularly following gene fusion events.

### Composite genes evolve multiple times independently

We quantified the rates at which evolution converged on the same composite gene using a phylogenetic approach, thus allowing us to overcome the bias related to gene loss. Using the phylogenetic signal within the homologous regions of the component and composite gene alignments, we determined the rate of independent origins of composite genes (**Figure 3**). We analysed component-composite gene trees to distinguish composite sequences that form monophyletic groups (which were most likely formed by a single event), from polyphyletic composite sequences (which represent possible multiple independent origins of that composite family, *e.g.*, multiple independent fusions/fissions). From among the 13,632 CHGs we selected families that met two criteria: whether they involved only two component families, and whether they contained more than three species in the alignment. In total, 10,829 of 13,632 CHGs satisfied these criteria.

Briefly, for each CHG we first aligned and trimmed the sequences and then built gene trees using IQTree (42, 43) (v2.03; using automatic model selection and carrying out 1000 ultrafast bootstrap replicates) for all homologous blocks of sequences of component and composite genes (**Figure 3a-b**). We then selected those maximum likelihood trees where the composite genes do not appear as a monophyletic group (9,124/10,829 or 84%). We re-ran the analysis using the same phylogenetic models as before, but this time we imposed topological constraints on the search for optimal trees, where we forced the composite sequences to form a monophyletic group (**Figure 3c**). The Approximately Unbiased (AU) test (44) was used to measure the significance in the difference in support for the unconstrained (polyphyletic) versus the constraint (monophyletic) tree (**Figure 3d**). The null hypothesis is that there is no significant difference in likelihood score for the constraint tree and the unconstrained tree. This approach provides a robust statistical framework to infer the rate of independent evolution of composite genes. Out of the 9,124 CHGs of putative independent origin, 5,631 rejected the null hypothesis, *i.e.* the monophyletic constraint is significantly worse than the polyphyletic gene tree implying independent origin of the same composite event. For the remaining 3,493 CHGs tested for single or multiple origins, there was no significant difference between the constraint and gene trees, *i.e.* a monophyletic origin could not be discounted and in these cases we parsimoniously assumed monophyletic origin. Whilst different joining-points are possible in both monophyletic and polyphyletic CHGs, there should be less constraint on identical joining-points in polyphyletic cases. To test this we annotated the joining-point in all composite genes for each CHG and asked whether this location fell within the same general region of the protein for each gene in a CHG (see Materials and methods). Indeed, an analysis of the joining-points of all CHGs shows that 70% of polyphyletic CHGs have different joining-points as compared to 44% for monophyletic CHGs. To summarise, across all 13,632 CHGs identified, 5,198 CHGs (38%) were most likely monophyletic, 5,631 CHGs (41%) emerge independently more than once across the animal phylogeny, and 2,803 CHGs (21%) could not be assessed in this way as a consequence of having more than two components and/or insufficient species in the alignment.

Next, we assessed whether there was a significant difference in the types of protein domains present in composite genes of multiple origin as compared to those of single origin. We found that domains with functions related to protein binding and binding (GO:0005515, GO:0005488) were enriched in the set of composite genes which emerged more than once (p<0.05, Benjamini-Hochberg correction) (**Supplementary Table S1**).

### Composite gene gain and loss events are concentrated at specific nodes on the phylogeny

In order to determine the tempo of CHG gain and loss across the animal phylogeny we analysed a subset of 10,829 (from a total of 13,632) CHGs where we could categorise the CHG as single or multiple origin. For the 5,198 CHGs of single origin we analysed their evolutionary history using the irreversible Dollo model of evolution (45) implemented in the R package Claddis (46, 47). For the 5,631 CHGs of multiple origin we used the reversible Mk model (48) implemented in the RevBayes software program (49). We used the species tree shown in **Figure 4** (50–52), the most appropriate model as described above, and applied time calibrations extracted from the TimeTree database (53). Using TimeTree as a source for divergence times allowed us to use a detailed phylogeny including all taxa of interest. However, TimeTree divergence times are summary statistics from a diversity of studies some of which are by now only of historical value. In particular, for the deep part of the animal phylogeny, TimeTree estimates are most likely too old (54). Accordingly, rates of origin and loss of composite genes estimated here should be interpreted as minimal estimated values, as reducing the branch length in the timetree following more recent animal divergence time studies will cause the inference of higher rates of origin and losses. We also employed an alternative rooting for the tree placing the Ctenophore as the earliest diverging group (55–57) and found no significant change to the results presented here (**Supplementary Figure S3**).

**Figure 4.**
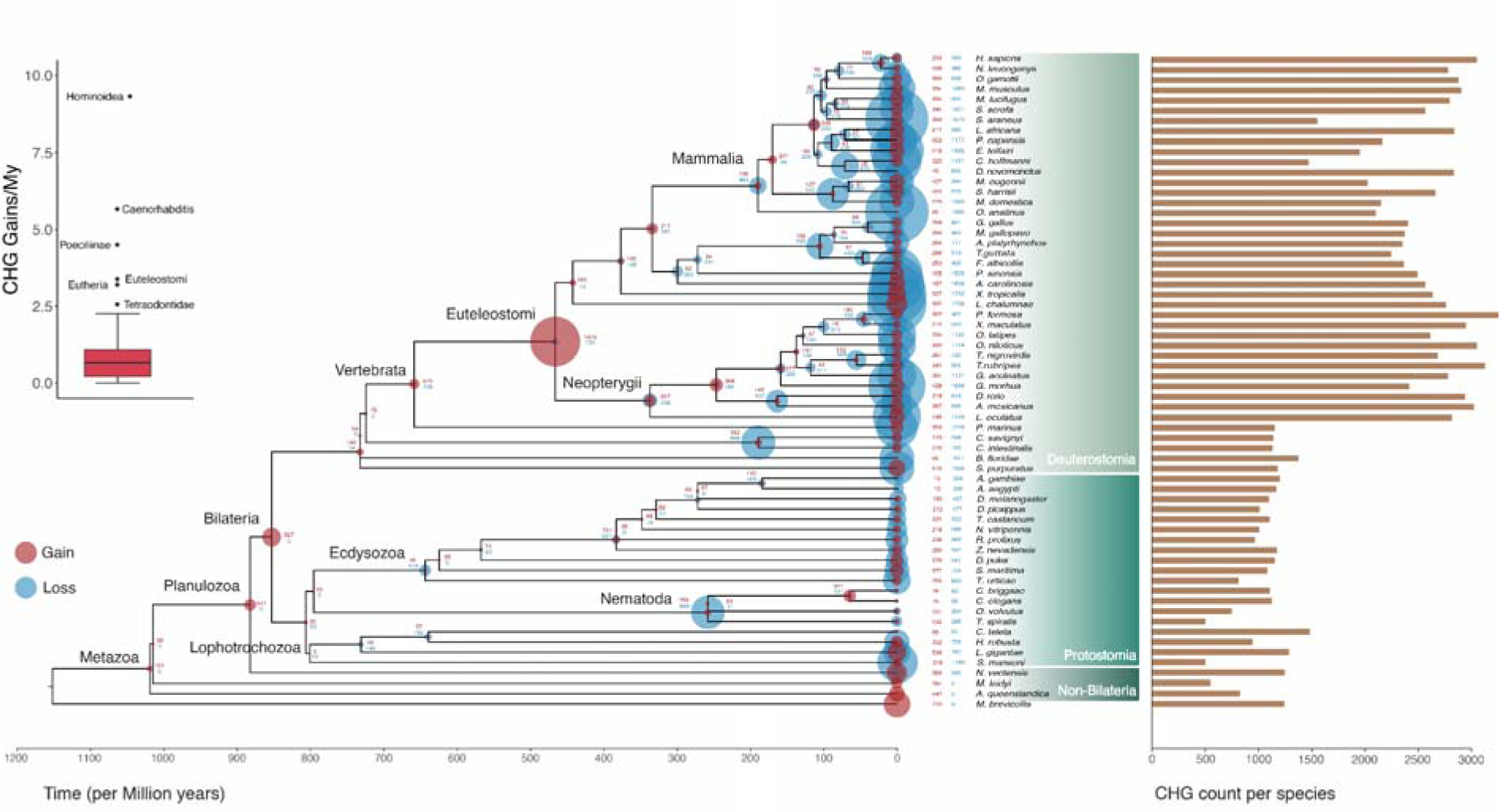
Distribution of the gain and loss of CHGs across the animal tree. The species phylogeny for our sample set is shown in the centre, with divergence time estimates in millions of years ago (MYA) on the x-axis (taken from TimeTree (53)). Gains of CHGs are shown as red discs and losses as blue. The size of the disc on the node is proportional to the amount of gain/loss at that node. The associated histogram on the right shows the total number of CHGs identifiable in the genomes of extant species. **Inset:** boxplot showing overall rate of gain for each node in the phylogeny, outlier nodes are named.

The first major observation is that CHG gain is not evenly distributed across the tree (**Figure 4**). Within Deuterostomia for example there are 21,381 separate CHG gain events across all branches compared to 6,295 CHGs gains in all branches within Protostomia. There are 1,322 gains in total across the five nodes preceding the divergence of Deuterostomia and Protostomia (*i.e.* Bilateria, Panulozoa, Eumetazoa, Metazoa, and Metazoa + Choanoflagellates). While there is a clear disparity between the number of composite genes between protostome and deuterostome species, this trend does not hold at the level of total gene count per species (**Supplementary Figure S4A&B**) suggesting underlying variation in the rate of CHG formation between these clades. To account for the difference in the number of species sampled between Protostomia and Deuterostomia, CHG counts for a random sample of 10 species were carried out 100 times, and we find that the number of CHGs present in the Deuterostome clade is significantly higher (Wilcoxon rank-sum test, *p*=0) (**Supplementary Figure S4C**). Finally, the distribution of CHGs across the tree suggests that a large proportion may be clade-specific. For example, there are 341, 338, and 1,475 CHGs unique to Caenorhabditis, Eutheria and Euteleostomi, respectively. The ancestral node with the largest number of CHGs gained was Eutelostomi with 1,475 gains, followed by Bilateria with 527 gains (**Figure 4**).

Across the tree we find that the rates of composite gene gain and loss per million years (MY) are highly variable and non-clock like (**Figure 4**). Note that the absolute values presented for rates are likely to be underestimated given the tree, but the comparisons of gains and losses remain valid. Within the internal branches of the phylogeny the average rate of CHG gain is 1.03/MY, compared to a loss rate of 2.81/MY. The branch leading to the Hominoidea clade displays the highest rate of CHG gain (9.31 CHG gains/MY). The branches leading to Caenorhabditis (5.67 gains/MY), Euteleostomi (3.39 gains/MY), Poecilinae (4.50 gains/MY), Eutheria (3.20 gains/MY), and Tetraodontidae (2.56 gains/MY) all display rates of CHG gain above the average plus the SD observed across the whole phylogeny (1.03 ± 1.51 CHG gains/MY) (**Figure 4 inset, Supplementary Figure S5A**). Contrastingly, higher rates of CHG loss tend to be found towards the tips of the tree, consistent with the observation of loss most often relating to loss of a single composite gene within a specific species rather than complete loss of the CHG (**Figure 4**). Branches with rates of loss higher than the average plus SD (2.81 ± 4.33 losses/MY) include: Hominoidea (24.96 losses/MY), Xenarthra (12.27 losses/MY), and Passeriformes (10.53 losses/MY), and the tip lineages: *Nomascus leucogenys* (23.08 losses/MY), *Choloepus hoffmanni* (21.98 losses/MY) (**Supplementary Figure S5B**). When assessing rates of gain and loss of composite genes, it must be noted that loss can relate to a complete loss of a composite gene from a species, or a reversal of the composite formation event (*e.g.* subsequent fission in a lineage following a fusion event). Gain of an individual composite gene member also occurs but at a lower rate than CHG member loss.

Finally, to assess the functional contribution of gene fusions specifically from our composite gene repertoire we compared the functional categories of fusion genes versus genes never associated with composite formation (*i.e*. neither composite nor components). We calculated the relative representation of Cluster of Orthologous Groups (COG) categories for genes of each type (*i.e.* fusion genes versus non-composite associated genes). We found that, in comparison to non-composite associated genes, fusion genes had a larger number of genes involved in transcription (K), post-translational modification, protein turnover, and chaperone functions (O), inorganic ion transport and metabolism (P), signal transduction (T), and extracellular structures (W) (**Supplementary Figure S6A**). To assess the potential functional impact of gene fusion events at major nodes in the animal tree, we measured the relative proportions of COG categories for fusion genes gained at these particular nodes (**Supplementary Figure S6B**). The overall proportions of COG categories are similar for all nodes tested, with signal transduction (T), transcription (K), and post-translational modification, protein turnover, and chaperone functions (O) representing the highest proportion of COG categories in all nodes. These patterns overlap with the broader contribution of fusion genes to these functions relative to non-composite genes, as seen above (**Supplementary Figure S6A**). Some clade specific shifts in COG proportions were observed; with overall larger proportion of genes involved in RNA processing and modification (A) present in the nodes Deuterostomia, Tetrapoda, Mammalia, and Eutheria; a larger contribution of extracellular structures (W) category in Euteleostomi and Tetrapoda; and a higher proportion of genes involved in defence mechanisms (V) in Tetrapoda, relative to other major nodes in the tree (**Supplementary Figure S6B**). Generally, the functional impact of gene fusion seems to be specific to certain broad functional categories throughout the animal phylogeny, pointing to a specific, <u>persistent</u>, role of gene fusion in driving the evolution of certain functions important for animal evolution (58).

## Discussion

The most recognisable part of evolutionary biology is the Tree of Life with its continually diverging branches emerging from the root. This narrative has hugely influenced how we think about evolutionary history, and it influences what we expect to see when we examine genomes. In addition to, and perhaps influenced by, the Tree of Life perspective, there is a feeling that evolution rarely, if ever, repeats itself. This last idea was most forcefully expressed by the palaeontologist Stephen Jay Gould who asked whether the tape of life was replayable (59) – a question to which Gould answered: No.

Fortunately, with the sequencing of an extensive array of genomes from many taxa across the diversity of animals we can address issues relating to the non-treelike aspects of evolution on one hand, as well as whether genome evolution is contingent on genetic background. If evolution is contingent on prior evolutionary events, then we expect that with increasingly divergent genetic backgrounds we are less likely to see repeated evolution, while on the other hand if evolution is largely deterministic, then despite differences in genetic backgrounds we expect to see the same evolutionary events occurring in different lineages. In the end, we find that the repeatability of evolution falls short of being hugely deterministic – only a small proportion of the overall animal gene repertoire shows evidence of repeated evolution, but by the same token, we see that repeatability has happened and it has happened across animal evolution many, many times.

Composite gene evolution is largely characterised by high rates of gene turnover, unequal rates of gain and loss between animal phyla, and significant bursts of composite gains intensified at particular nodes in the animal phylogeny. The rate of composite gene formation varies drastically between the major animal groups, with a greater number of composite genes found in Deuterostomia, compared to Protostomia or the non-bilaterian lineages. The single largest number of CHG gain events observed are on the branch leading to the Euteleostomi ancestor - a branch synonymous with major phenotypic innovations such as the emergence of mineralised bone and the development of a more complex immune system. The higher rates of CHG birth on this branch may also be related to increases in genome complexity at this point in animal evolution (60, 61). Multiple whole genome duplication (WGD) events within the vertebrate clade (62), coincide with nodes containing a large number of composite genes. In terms of rate of composite gain per million years, the branch leading to Euteleostomi also shows a significantly higher rate of gain than the average across the whole tree, reinforcing the contribution of gene fusion and fission at this point in animal evolution. The branch leading to the Caenorhabditis species also shows significantly higher rates of CHG gain. While this branch may not represent a point of major phenotypic or genomic change, species of this phylum are known to have a higher rate of recombination within their genomes (63). This may provide a molecular basis for the increased rates of gene fusion and fission in this group. Compared to patterns of gain and loss of other gene types, which show significant gain in early metazoan branches and pre-metazoan branches in particular (17, 18, 20), we find that the highest rates of composite gene gain correlate with nodes that emerge subsequent to these deep nodes, suggesting that the evolution of genetic content through mechanisms other than fusion and fission may be followed by subsequent higher rates of composite gene formation.

In animals, convergent fusion events are known, for example the TRIM5-CYPA gene in New World monkeys (64) and Old World monkeys (65), the repeated fusion of β-globin genes in Laurasiatheria (66), and the recurrent fusion of transcription factors and transposons in vertebrates (67). More broadly speaking, 25% of all multi-domain proteins in eukaryotes are thought to have emerged independently, and 71% of domain combinations in the human genome have been found to be gained independently in at least one other eukaryotic genome (27). There have also been several examples of recurrent gene fusion in different eukaryote lineages (68, 69). Our estimate for the rate of convergent evolution of composite genes in the evolution of animals suggests that selection for the same combinations of gene sequences in composite genes is indeed common, with 41% (5,631 CHGs) having likely evolved independently more than once on the tree. Given our approach, using phylogenetic signal within the composite and component sequences, we could be confident that our results are not skewed by taxon sampling or data quality issues. The data presented here suggests that there are high levels of CHG formation and loss across animal evolution, that the same composites form independently across the tree, and that these CHG likely contribute substantially to the emergence of animal gene repertoires providing functional innovation, *e.g.* the IGFBP fusion protein. This work has important implications for our understanding of how protein coding genes evolve in animals, the prevalence of convergent evolution, how we construct gene families, and how we annotate function between homologous genes.

## Author Contributions

MJO’C and JMcI conceived of the study. MJO’C, JMcI, EB, PL, CC, RM, JP, and PM designed the specific experiments. PM, RM, and JP performed most experiments. RM, JP, PM, MJO’C, EB, JMcI, and PL designed, performed, and interpreted the homology and composite searching analyses. DS, EP and PM performed functional enrichment analyses, and DS, EB and PL performed the fission vs fusion analyses. PM, RM, JP, CC, KST, BC, PL and FJW contributed code for data analyses and data visualisation. MJO’C, PM, RM, JP, JMcI, DP, KST, DP and CC contributed to the interpretation of the results. PM, RM, JMcI and MJO’C took the lead in writing the manuscript. All authors provided critical feedback and helped shape the overall manuscript.

## Classification

Biological Sciences, Evolution.

## Data availability

All data and code used in this study are publicly available. Find all necessary information deposited at: https://figshare.com/projects/CompositeGenesMetazoa_Mulhair_et_al_/127943

## Acknowledgements

This work was undertaken on ARC3, part of the High-Performance Computing facilities at the University of Leeds, UK. We thank Martin Callaghan and all members of the ARC team for their excellent technical support. PM was funded through a University Academic Fellowship to MOC at the University of Leeds. RM was funded by the IRCSET PhD scholarship (GOIPG/2014/306). The authors wish to acknowledge the Irish Centre for High-End computing (ICHEC) for the provision of computational facilities and support. EB and JP were supported by a FP7/2007-2013 Grant Agreement # 615274, category LS8). C.J.C. wishes to acknowledge funding from the European Commission via Horizon 2020 (818368, MASTER with K.S.T. and 101000213 HoloRuminant); FJW was supported by a Marie Skłodowska-Curie Individual Fellowship (GA no. 793818) and a University of Nottingham Anne McLaren Fellowship. DP wishes to acknowledge funding from the John Templeton Foundation (#62220 although the opinions expressed in this paper are those of the authors and not those of the John Templeton Foundation) and the Gordon and Betty Moore Foundation (GBMF9741). We would also like to thank all current and former members of the O’Connell and McInerney research groups for invaluable discussions and insights over the course of this work. We are publishing under an open access license.

