## Supplementary Files (Figures and Tables) for "Bursts of novel composite gene families at major nodes in animal evolution"

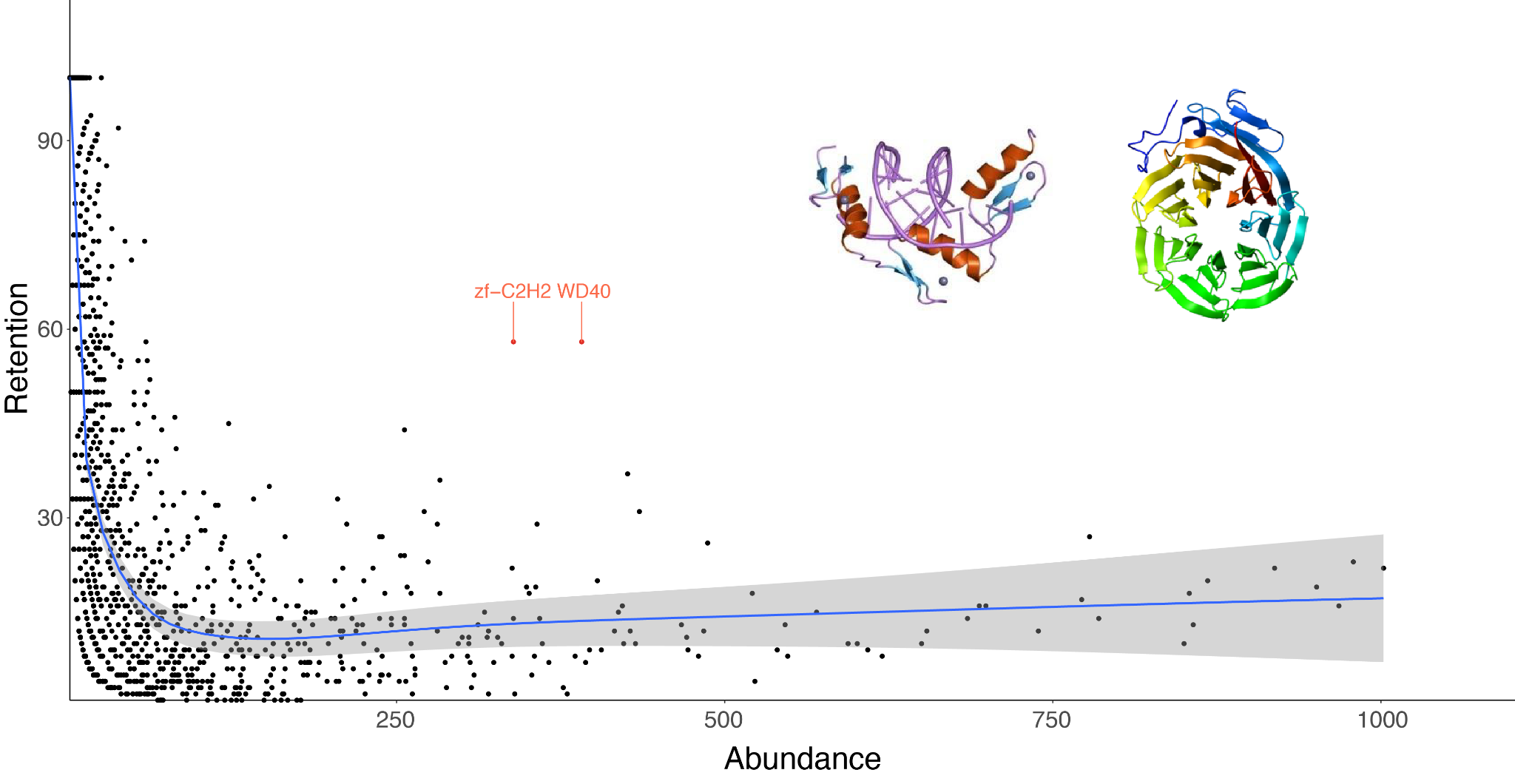

***Supplementary Figure S1.*** ***Rate of retention of domains in composite formation events.*** Each point represents a protein domain as annotated through Pfam. The domains are plotted by the number of times they are present in a component gene family (Abundance) by the percentage of time they are passed onto the composite gene from the component gene (Retention). Highlighted are the two most outlier cases, zf-C2H2 and WD40, which are present and retained at a higher rate than any other domains.

***Supplementary Figure S2.*** ***Contribution and placement of fusion versus fission to composite gene formation.*** **(A)** Species phylogeny (left) with species names coloured depending on whether they are non-bilaterian (black), present in Protostomia (dark green), or Deuterostomia (light green). Stacked bar chart shows the proportion of genes in each species annotated as non-composite associated (i.e. neither composite nor component), a fusion gene, or a fission gene. **(B)** Species phylogeny with fusion genes mapped onto their node of origin, as determined by parsimony placement used to distinguish between whether a composite gene is a fusion or a fission (with respect to where the component genes place on the tree). **(C)** Species phylogeny with fission gene events mapped onto their node of origin. In both **(B)** and **(C)**, species tips names are coloured as in **(A)** to distinguish between non-Biltaria, Protostomia, and Deuterostomia.
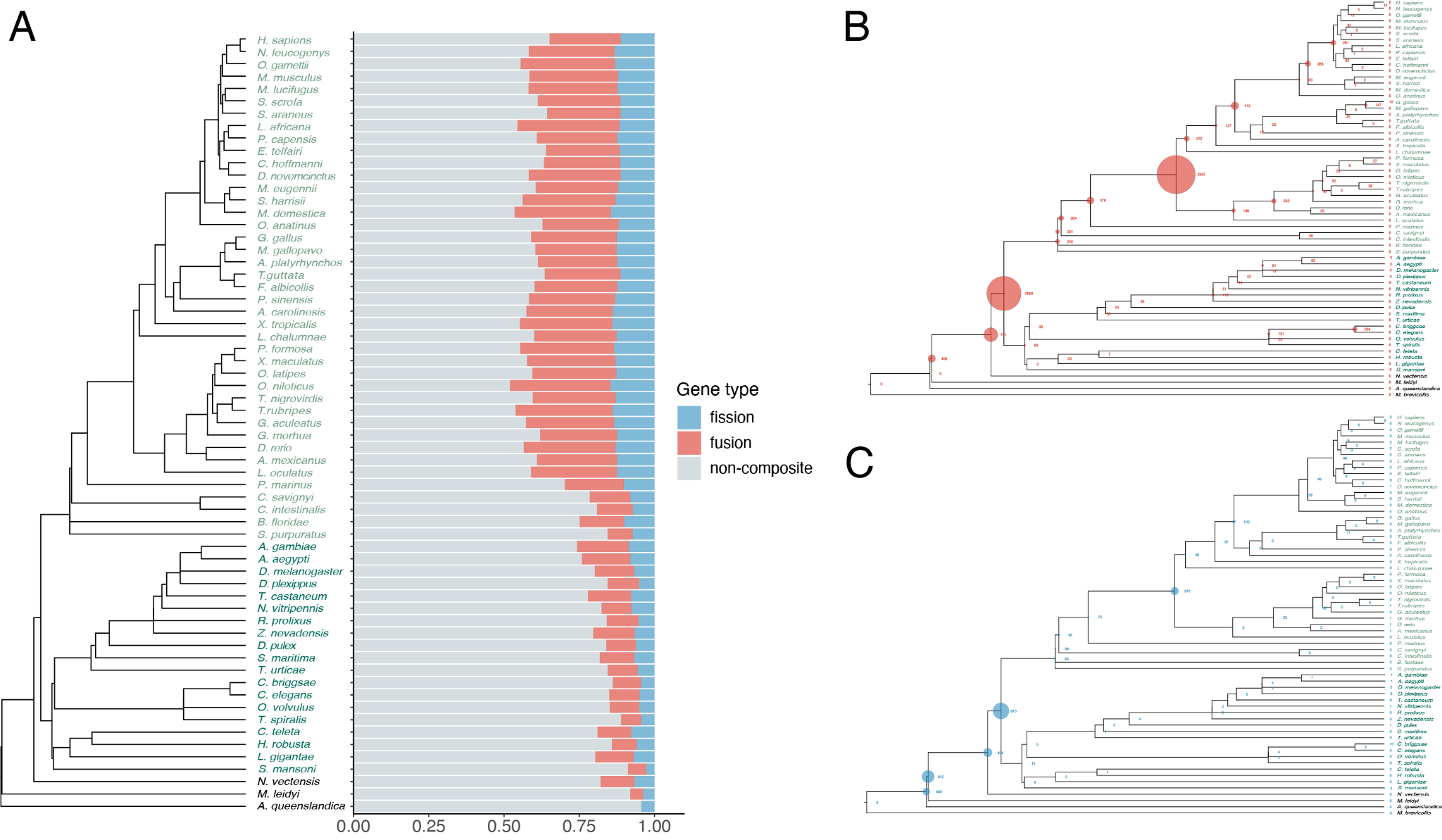

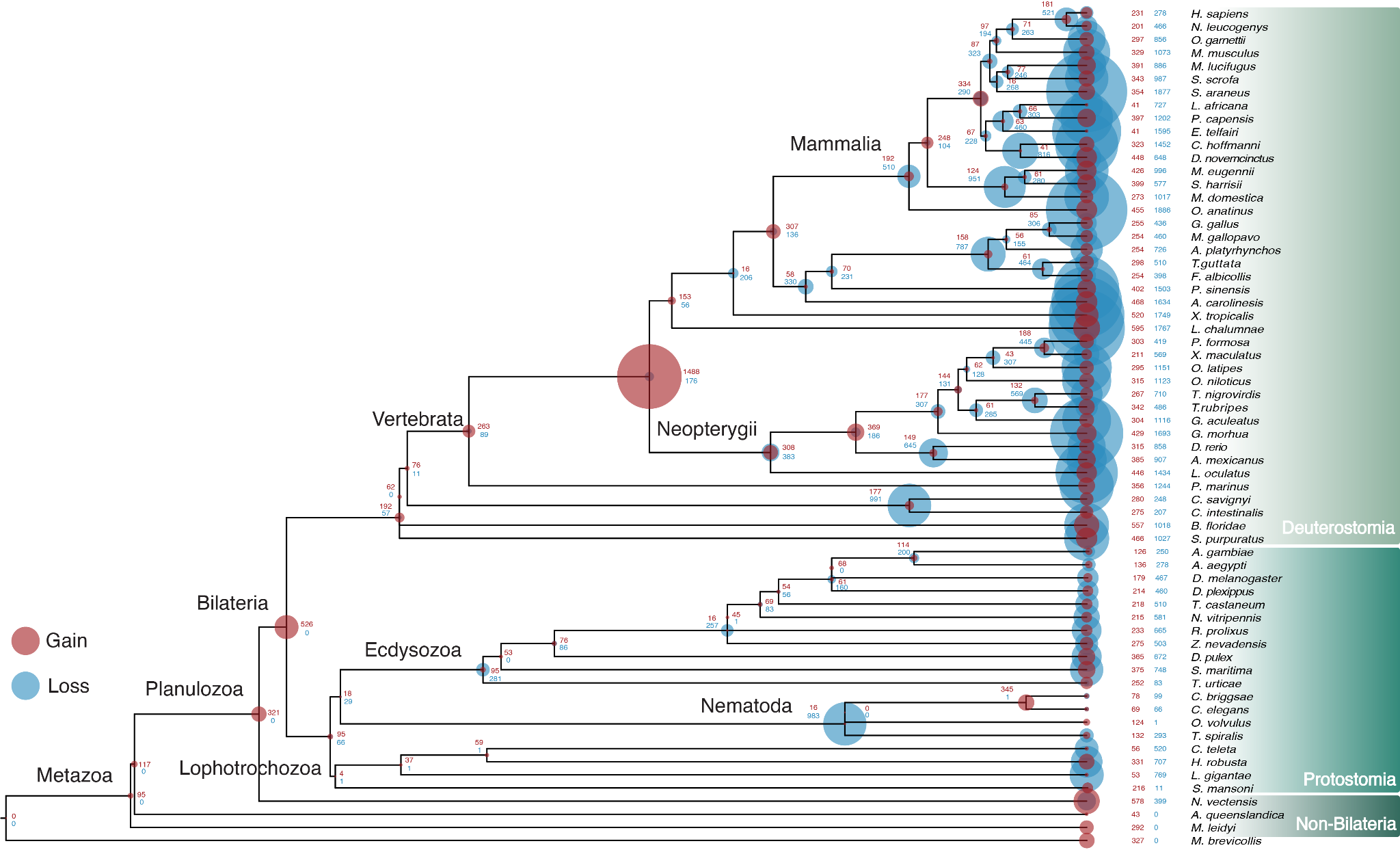

***Supplementary Figure S3. Patterns of CHG gain and loss event across “Ctenophore-sister” rooting of the animal tree.*** The species phylogeny here places ctenophores as the earliest branching animal in our phylogeny. As per **Figure 2**, again the gains of CHGs are shown as red discs and losses in blue and the size of the disc on the node is proportional to the amount of gain/loss at that node.

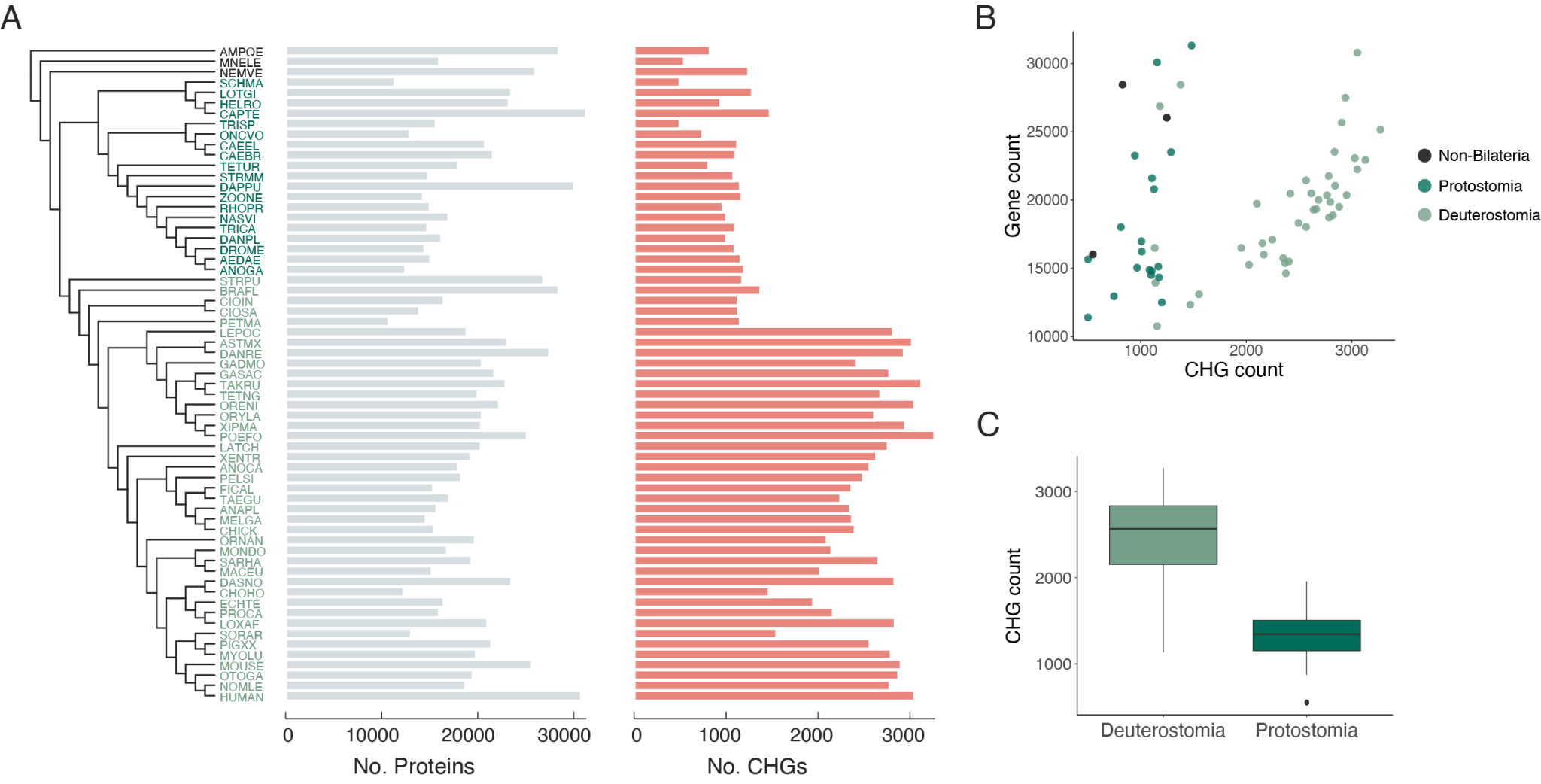

***Supplementary Figure S4. Comparison of CHG number between Protostomia and Deuterostomia.* (A)** Phylogeny of 63 animal species with tips coloured based on whether species are non-bilaterians, protostomes, or deuterostomes. Bar charts show the total number of proteins (left, grey) and composite proteins (right, red) per species. **(B)** The number of composite genes (x-axis) in relation to the total number of genes (y-axis) in each of the three major clades. **(C)** The distribution of CHG counts for a random sample of 10 Deuterostome/Protostome species repeated 100 times, each random sampling is shown (Wilcoxon rank-sum test, *p*=0).

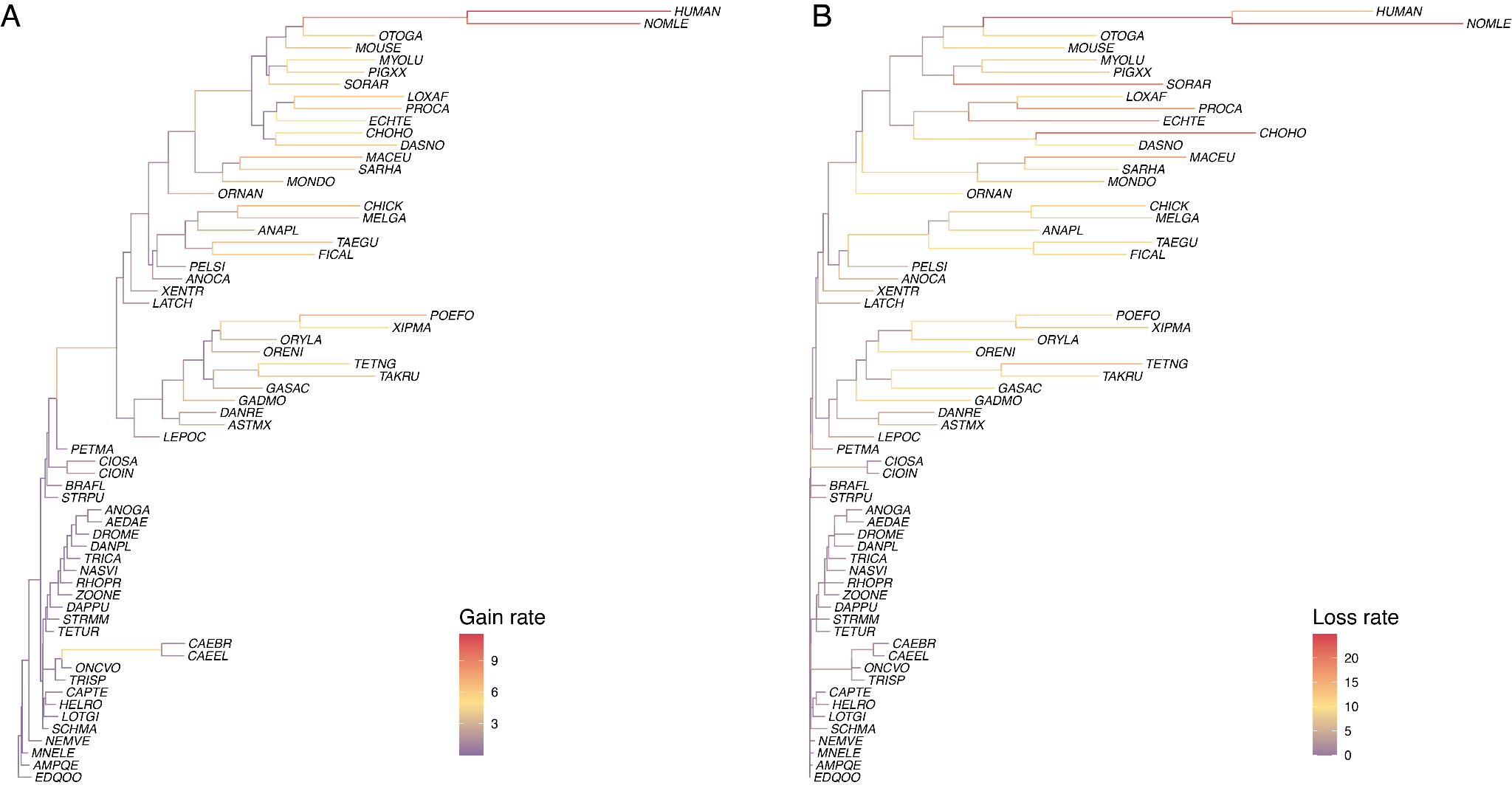

***Supplementary Figure S5 Rates of gain and loss of CHGs across the animal phylogeny.*** Branch lengths represent the number of CHGs **(A)** gained or **(B)** lost per million years. The branches are coloured along a gradient from purple representing 0, to red representing highest rates of gain or loss. For **(A)** the scale of gains of CHGs/MY ranges from 0 (purple) to 12 (red) and **(B)** the scale of losses of CHGs/MY ranges from 0 (purple) to 25 (red).

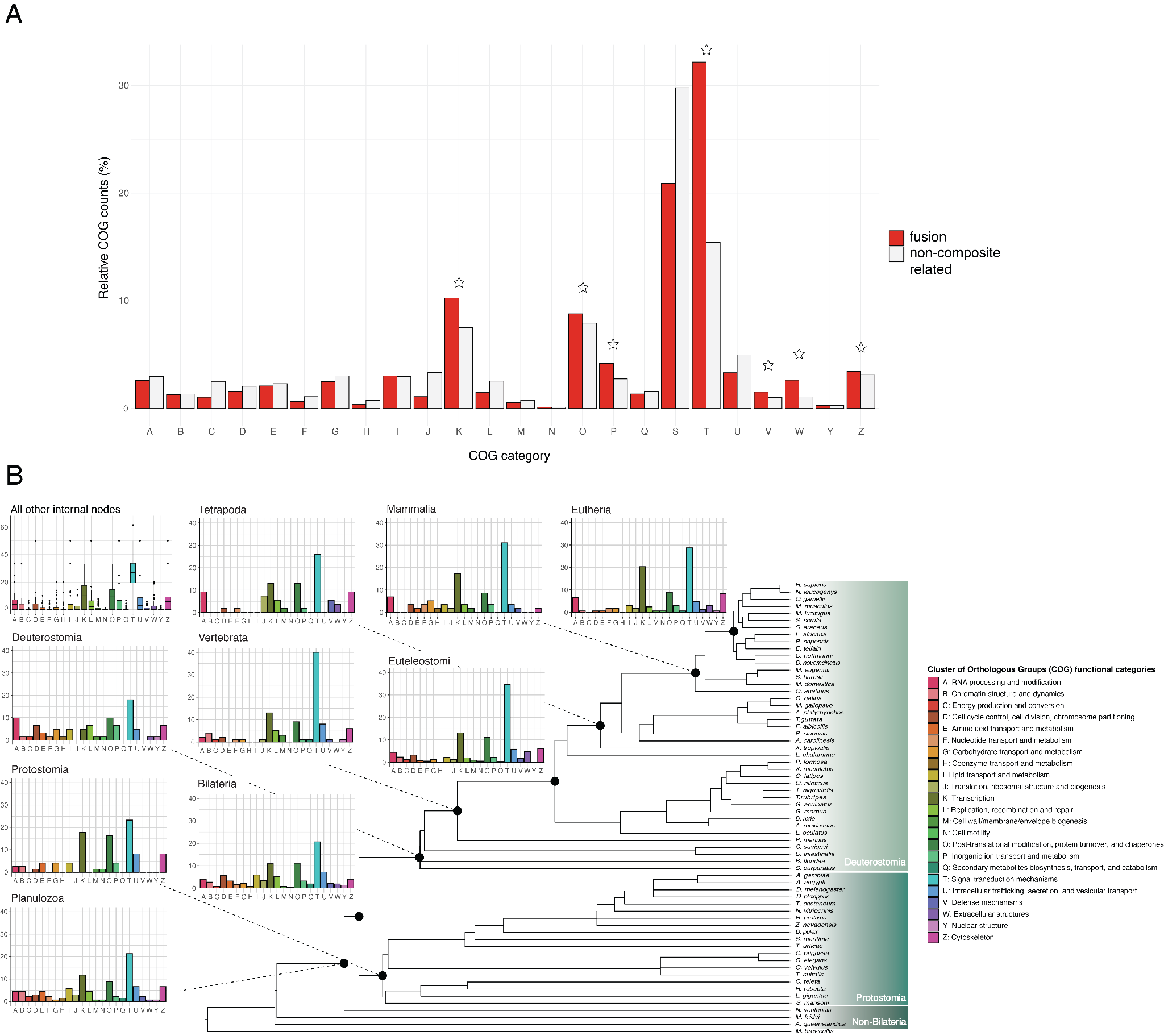

***Supplementary Figure S6: Comparison of COG functional categories across gene types and across the tree.* (A)** Histogram representing the proportions of fusion genes (red) or non-composite associated genes (grey) (*y-axis*) annotated with each of the COG functional categories (*x-axis*). Single letter codes for functional categories are shared with panel B below. Stars signify categories where the proportion of fusion genes is higher than that of non-composite associated genes. **(B)** Animal phylogeny with proportions of COG categories highlighted for all CHGs gained at (i) major internal nodes (annotated with the clade name above), and (ii) the overall proportions of COG categories for CHGs gained at all other nodes in the tree (top left). COG functional categories are listed on the right with their associated single letter code and their colour.

| **GO** | **NS** | **enrichment** | **name** | **ratio_in_study** | **ratio_in_pop** | **p_uncorrected** | **depth** | **study_count** | **p_fdr_bh** | **study_items** |
| --- | --- | --- | --- | --- | --- | --- | --- | --- | --- | --- |
| GO:0005515 | MF | e | protein binding | 61/583 | 164/3442 | 0.000000000152242553 | 2 | 61 | 0.0000001032204509 | 4_1_CTD, ASD1, Ank, BTB, Bromodomain, CAMSAP_CC1, CUE, CaMKII_AD, DED, Death, Dimer_Tnp_hAT, E3_UbLigase_EDD, EF-hand_4, EMI, F-box, F-box-like, FATC, FOLN, Fz, GYF, HLH, IGFBP, IQ, Kazal_2, Kelch_1, LRR_6, LRR_8, Ldl_recept_a, NHL, PAS_3, PAZ, PDGF, PDZ, PLAT, RWD, Rapsyn_N, SAM_1, SAM_2, SET, SH3_1, SH3_2, SH3_9, SPRY, SRF-TF, SWIRM, Sema, Spectrin, TGF_beta, TIR, TNFR_c6, TPR_1, TPR_8, Ubiquitin_2, VHP, VWC, W2, WD40, WH2, WW, fn3, ubiquitin |
| GO:0005488 | MF | e | binding | 136/583 | 545/3442 | 0.0000002011833857 | 1 | 136 | 0.00006820116775 | 3HCDH_N, 4_1_CTD, 5-nucleotidase, ABC_membrane, ABC_tran, ACBP, AIG1, ARID, ASD1, Acyl-CoA_dh_N, Alpha_kinase, Androgen_recep, Ank, Arf, BTB, Bromodomain, CAMSAP_CC1, CUE, CaMKII_AD, Cadherin, Cpn60_TCP1, DDE_1, DEAD, DED, DM, Death, Dimer_Tnp_hAT, E3_UbLigase_EDD, EF-hand_1, EF-hand_4, EF-hand_6, EF-hand_7, EGF_CA, EMI, Ets, F-box, F-box-like, FATC, FMO-like, FOLN, FYVE, Fz, G-alpha, G-patch, GATA, GTP_EFTU_D2, GYF, Gal-bind_lectin, Gal_Lectin, HIRAN, HLH, HMA, HNOB, HSP70, Hairy_orange, IGFBP, IQ, KH_1, Kazal_2, Kelch_1, LBP_BPI_CETP, LBP_BPI_CETP_C, LRR_6, LRR_8, Ldl_recept_a, Linker_histone, MCM, MMR_HSR1, Myosin_head, NHL, PABP, PAS_3, PAZ, PDGF, PDZ, PLAT, PX, Peptidase_M10, Pkinase, Pre-SET, RINGv, RNase_H, RRM_1, RWD, Rapsyn_N, Ras, Rep-A_N, SAM_1, SAM_2, SAM_PNT, SET, SH3_1, SH3_2, SH3_9, SNF2_N, SPARC_Ca_bdg, SPRY, SRF-TF, SRP54, SRP54_N, SRP_SPB, SWIRM, Sema, Septin, Sod_Cu, Spectrin, TGF_beta, TIR, TNFR_c6, TPR_1, TPR_8, Ubiquitin_2, VHP, VWC, W2, WD40, WH2, WW, Xlink, eIF-5_eIF-2B, fn3, p450, tRNA-synt_2, tRNA_anti-codon, ubiquitin, zf-AD, zf-AN1, zf-B_box, zf-C2HC, zf-C3HC4, zf-C4, zf-CCHC, zf-CXXC, zf-U1, zf-UBP, zf-UBR |
| GO:0003674 | MF | e | molecular_function | 220/583 | 1005/3442 | 0.000001165592771 | 0 | 220 | 0.0002634239663 | 3HCDH, 3HCDH_N, 4_1_CTD, 5-nucleotidase, 7tm_1, 7tm_2, 7tm_3, A2M, AA_permease_2, ABC_membrane, ABC_tran, ACBP, AIG1, ARID, ASD1, ATP-gua_Ptrans, ATP-gua_PtransN, Abhydrolase_3, Acetyltransf_1, Acyl-CoA_dh_1, Acyl-CoA_dh_M, Acyl-CoA_dh_N, Acyltransferase, Alpha-amylase, Alpha_kinase, Amino_oxidase, Androgen_recep, Ank, Antistasin, Arf, ArfGap, Asparaginase_2, Astacin, B56, BIRC6, BTB, Bin3, Bromodomain, CAMSAP_CC1, CUE, CaMKII_AD, Cadherin, Cpn60_TCP1, DAGK_acc, DAGK_cat, DAO, DDE_1, DEAD, DED, DHHC, DM, Death, Dimer_Tnp_hAT, E3_UbLigase_EDD, ECH_1, EF-hand_1, EF-hand_4, EF-hand_6, EF-hand_7, EGF_CA, EMI, Ets, F-box, F-box-like, FAD_binding_8, FATC, FKBP_C, FMO-like, FOLN, FYVE, Formyl_trans_N, Fz, G-alpha, G-patch, GARS_C, GARS_N, GATA, GNT-I, GTP_EFTU_D2, GYF, Gal-bind_lectin, Gal_Lectin, Glutaredoxin, Glyco_hydro_2, Glyco_hydro_2_C, Glyco_hydro_2_N, Glyco_transf_10, Glyco_transf_29, GoLoco, HECT, HIRAN, HLH, HMA, HNOB, HNOBA, HSL_N, HSP70, Hairy_orange, Hydantoinase_A, Hydantoinase_B, IGFBP, IQ, ITI_HC_C, Ion_trans, KH_1, Kazal_2, Kelch_1, Kunitz_BPTI, LBP_BPI_CETP, LBP_BPI_CETP_C, LRR_6, LRR_8, Latrophilin, Ldl_recept_a, Lig_chan, Lig_chan-Glu_bd, Linker_histone, MCM, MFS_1, MMR_HSR1, MOZ_SAS, Med15, Myosin_head, NAD_binding_6, NCD3G, NHL, Na_H_Exchanger, Na_sulph_symp, PABP, PAS_3, PAZ, PDGF, PDZ, PIP5K, PLAT, PX, Peptidase_C13, Peptidase_M10, Peptidase_S10, Phospholip_A2_1, Pkinase, Pre-SET, Pro_isomerase, RGS-like, RINGv, RNase_H, RRM_1, RWD, Rapsyn_N, Ras, Rep-A_N, Reprolysin, RhoGEF, Ribonuclease_3, Ribosomal_L19e, Ribosomal_L44, SAM_1, SAM_2, SAM_PNT, SDF, SET, SH3_1, SH3_2, SH3_9, SLC12, SMG1, SNF2_N, SPARC_Ca_bdg, SPRY, SRF-TF, SRI, SRP54, SRP54_N, SRP_SPB, SSF, SWIRM, Sema, Septin, Sod_Cu, Spectrin, Sulfatase, Sulfotransfer_1, T4_deiodinase, TGF_beta, TIR, TNFR_c6, TPR_1, TPR_8, Tme5_EGF_like, Trypsin, UCH, UDPGT, Ubiquitin_2, VHP, VWC, W2, WAP, WD40, WH2, WW, Xlink, Y_phosphatase, bZIP_1, eIF-5_eIF-2B, fn3, p450, tRNA-synt_2, tRNA_anti-codon, ubiquitin, zf-AD, zf-AN1, zf-B_box, zf-C2HC, zf-C3HC4, zf-C4, zf-CCHC, zf-CXXC, zf-U1, zf-UBP, zf-UBR |

***Supplementary Table S1. List of enriched GO terms associated with pfam domains present in composite genes which emerged multiple times independently.*** List of GO terms enriched in composite genes which emerged convergently (following analysis in Figure 3). Goatools find_enrichment.py script was used to assess enrichment in convergently evolved composite genes relative to the background of all composite genes in the dataset.

| **Original species** | **Replacement species** |
| --- | --- |
| *Astyanax mexicanus* | *Hydrolycus armatus* |
| *Lottia gigantea* | *Lottia kogamogai* |
| *Oreochromis niloticus* | *Oreochromis tanganicae* |
| *Poecilia formosa* | *Poecilia reticulata* |
| *Strongylocentrotus purpuratus* | *Mesocentrotus franciscanus* |
| *Nasonia vitripennis* | *Nasonia giraulti* |
| *Onchocerca volvulus* | *Brugia malayi* |
| *Schistosoma mansoni* | *Schistosoma sinensium* |
| *Tetranychus urticae* | *Gozmanyina majesta* |
| *Trichinella spiralis* | *Ascaris suum* |

***Supplementary Table S2. Replacement species used to place divergence time for species in our dataset.*** For the twelve species in our dataset missing from the TimeTree database (left column), closely related species to these lineages were used as replacements (right column).
